# Spinal stretch reflexes support efficient control of reaching

**DOI:** 10.1101/2020.01.06.896225

**Authors:** Jeffrey Weiler, Paul L Gribble, J. Andrew Pruszynski

## Abstract

Efficiently controlling the movement of our hand requires coordinating the motion of multiple joints of the arm. Although it is widely assumed that this type of efficient control is implemented by processing that occurs in the cerebral cortex and brain stem, recent work has shown that spinal circuits can generate efficient motor output that supports keeping the hand in a static location. Here, we show that a spinal pathway can also efficiently control the hand during reaching. In our first experiment we applied multi-joint mechanical perturbations to participant’s elbow and wrist as they began reaching towards a target. We found that spinal stretch reflexes evoked in elbow muscles were not proportional to how much the elbow muscles were stretched but instead were efficiently scaled to the hand’s distance from the target. In our second experiment we applied the same elbow and wrist perturbations but had participants change how they grasped the manipulandum, diametrically altering how the same wrist perturbation moved the hand relative to the reach target. We found that changing the arm’s orientation diametrically altered how spinal reflexes in the elbow muscles were evoked, and in such a way that were again efficiently scaled to the hand’s distance from the target. These findings demonstrate that spinal circuits can help efficiently control the hand during dynamic reaching actions, and show that efficient and flexible motor control is not exclusively dependent on processing that occurs within supraspinal regions of the nervous system.

## Introduction

Goal-directed movements are plagued by noisy motor signals (Faisal et al., 2008) and influenced by environmental forces. The simplest way to deal with these disturbances is to monitor sensory information from individual joints and locally correct any deviations away from each joint’s intended movement trajectory. However, the redundancy of the musculoskeletal system affords an abundance of motor solutions to compensate for such disturbances (Bernstein, 1967; Latash, 2000). Therefore, a more sophisticated and efficient control scheme is to ignore disturbances at the individual joint-level and instead only make corrective responses to the degree task success is hindered (Todorov, 2004; Todorov and Jordan, 2002).

There are numerous examples across a wide range of tasks showing that individuals generate efficient corrective responses (Cole et al., 1984; Cole and Abbs, 1987; Diedrichsen, 2007; Dimitriou et al., 2012; Gracco and Abbs, 1985; Mutha and Sainburg, 2009; Omrani et al., 2013; Robertson and Miall, 1997). For example, when moving a (virtual) object with two hands, disturbing the movement by mechanically perturbing one arm causes corrective responses at both arms (Diedrichsen, 2007; Mutha and Sainburg, 2009). These efficient corrective responses have been attributed to processing that occurs exclusively across supraspinal pathways (Pruszynski and Scott, 2012; Scott, 2016) because spinal pathways typically generate corrective responses at the joint-level proportional to how much that joint was disturbed (Pierrot-Deseilligny and Burke, 2012), consistent with the simplest manner of dealing with internal or external disturbances. However, we have recently shown that a spinal pathway is capable of exploiting the arm’s musculoskeletal redundancy to efficiently support postural control of the hand (Weiler et al., 2019). In that work we applied small mechanical perturbations to participants’ elbow and wrist while they maintained their hand at a home location. Critically, the perturbations that displaced the hand furthest from the home location did so with the least amount of elbow rotation (and vice versa). We found that spinal stretch reflexes were elicited in elbow muscles, but their magnitudes were not proportional to how much the elbow was rotated. Instead, these spinal reflexes were scaled in such a way to re-establish the hand’s position back at the home location – a result showing that a spinal pathway can concurrently process sensory information from multiple joints and generate efficient motor output to keep the hand in a stationary position.

There are important differences in how the central nervous system processes sensory information during posture compared to movement. For example, the amplitude of the H-reflex is relatively consistent during quiet stance but is dynamically modulated during movement (Capaday and Stein, 1986), and conscious percepts of somatosensory information can be reduced when moving compared to remaining stationary (Chapman et al., 1987; Duysens et al., 1990). But perhaps the most extensively documented difference is that sensory information from the periphery can be ‘gated’ during movement, leading to a marked reduction in afferent evoked activity in the spinal cord (Hantman and Jessell, 2010; Seki and Fetz, 2012), cuneate nucleus (Ghez and Pisa, 1972), thalamus (Tsumoto et al., 1975) and cortex (Papakostopoulos et al., 1975; Seki and Fetz, 2012). Given these differences, spinally-generated corrective responses that efficiently control the hand may be restricted to when the hand is stationary.

Here we tested whether a spinal pathway can support efficient hand control while reaching. In our first experiment we applied multi-joint mechanical perturbations to participants elbow and wrist once they began reaching to a target. In our second experiment we applied the same mechanical perturbations to the elbow and wrist but also had participants change their arm orientation, which diametrically altered how the same rotation of the wrist moved the hand relative to the reaching target. Critically, across both experiments, the multi-joint perturbations that moved the hand furthest from the target did so with the least amount of elbow rotation (and vice versa). We found that the spinal reflexes evoked in the elbow muscles were not proportional to how much the elbow was rotated – and thus how much the elbow muscles were stretched – but instead were efficiently scaled to the hand’s distance from the target. This finding demonstrates that the spinal pathway that processes feedback from multiple joints to efficiently control the hand is engaged in both static and dynamic contexts and shows that efficient and flexible motor control is not exclusively dependent on processing that occurs within supraspinal regions of the nervous system.

## Materials and Methods

### Participants

Twenty individuals volunteered to participate in *Experiment 1* (males = 6, females = 14, age range 19-23) and 12 individuals volunteered to participate in *Experiment 2* (males = 8, females = 4, age range = 19-38). All participants reported being free from neurological and musculoskeletal dysfunction and provided informed written consent prior to data collection. This work was approved by the Office of Research Ethics at Western University and was completed in accordance with the Declaration of Helsinki.

### Apparatus

Participants grasped the handle of a three degree-of-freedom exoskeleton robot (Interactive Motion Technologies), which allows movement of the hand in a horizontal plane by flexing or extending the shoulder, elbow and wrist (Weiler et al., 2015). The exoskeleton is equipped with direct-drive motors (rise-time = 2ms) that can independently apply flexion or extension torques at each of the aforementioned joint segments, and with 16-bit rotary encoders (Gurley Precision Instruments) to measure joint kinematics (resolution = 0.0055 degrees). Visual stimuli were presented by a 46-inch TV monitor onto a semi-silvered mirror that occluded vision of the participant’s arm. Participants were comfortably seated in a height adjustable chair and the lights in the experimental suite were extinguished during data collection.

### General Procedures

Participants were provided visual feedback of their hand position with a 1cm in diameter turquoise circle, which was displayed at the coordinates of the exoskeleton’s handle (i.e., hand feedback cursor). Each trial began with the participant moving the hand feedback cursor to a 1cm in diameter orange circle (i.e., start location), which corresponded to their hand’s location when the shoulder, elbow and wrist were at 80°, 70° and 10° of flexion, respectively (external angle coordinate system). After remaining at this location for 500ms, the exoskeleton gradually applied flexion torque at the elbow and the wrist for 1000ms that plateaued at 2Nm and 0.8Nm, respectively (i.e., the background load). A 1cm in diameter red circle was concurrently presented at this time (i.e., home location), which corresponded to their hand’s location when the shoulder, elbow and wrist were at 70°, 60° and 10° of flexion, respectively, and participants moved their hand to its location. When the participant moved the cursor to the home location and had their wrist between 5 and 15° of flexion, the home location changed from red to orange and the hand feedback cursor disappeared. After maintaining the position for a randomized foreperiod (i.e., 1000-2000ms), another 1cm diameter orange circle was presented (i.e., the target) and participants were instructed to move to its location when ready. This reaching movement required 10° of pure elbow extension. On half of the trials (see below) the moment the hand left the home-location the exoskeleton applied a 2Nm flexion step-torque (i.e., the perturbation) at the elbow and could simultaneously apply a ±0.8Nm flexion or extension step-torque at the wrist. The hand feedback cursor was re-shown 100ms after movement onset. If participants reached the target in <600ms, the target changed from orange to green; otherwise the target changed from orange to red. The loads applied by the robot were then quickly ramped down, which denoted the end of the trial.

### Experiment Specific Procedures

In Experiment 1, participants reached for the target in a single block of trials. At movement onset the robotic exoskeleton (1) mechanically flexed their elbow and flexed their wrist, (2) mechanically flexed their elbow and extended their wrist, (3) only mechanically flexed their elbow, or (4) applied no mechanical perturbations. Participants completed 60 trials for each of the three experimental conditions that contained a mechanical perturbation and 180 trials of the experimental condition in which no mechanical perturbation was applied in a randomized order, totalling 360 trials.

In Experiment 2, participants reached for the target in two separate blocks of trials. Critically, the blocks differed by how participants grasped the robot handle, which diametrically altered how wrist rotation moved the hand relative to the target. In one block, participants grasped the robot handle normally with their thumb pointing upward (i.e., Upright) whereas in the other block participants internally rotated their forearm and grasped the handle with their thumb pointing downward (i.e., Flipped). In both blocks of trials, at movement onset the robotic exoskeleton (1) mechanically flexed their elbow and flexed their wrist, (2) mechanically flexed their elbow and extended their wrist or (3) applied no mechanical perturbations. In both blocks of trials, participants completed 60 trials for each of the two experimental conditions that contained a mechanical perturbation and 120 trials for the experimental condition in which no mechanical perturbation was applied in a randomized order, totalling 480 trials. The ordering of the blocks was randomized across participants.

### Muscle Activity

Participants’ skin was cleaned with rubbing alcohol and EMG surface electrodes (Delsys Bagnoli-8 system with DE-2.1 sensors, Boston, MA) were coated with a conductive gel. Electrodes were then secured on the skin over the belly of the lateral head of the triceps brachii (TRI) at an orientation that runs parallel to the muscle fibers. A reference electrode was secured over the participants’ left clavicle. EMG signals were amplified (gain = 10^3^) and then digitally sampled at 2,000 Hz.

### Data Reduction and Statistical Analyses

All data were aligned to the moment the hand left the home location (i.e., movement onset), which also corresponded to when perturbations could be applied. Angular position of the shoulder, elbow and wrist were sampled at 500 Hz and low-pass filtered (12 Hz, 2-pass 2^nd^-order Butterworth). Participants hand position was computed based on the lengths of the robotic exoskeleton’s adjustable upper-arm, lower-arm and wrist linkages. EMG data were band-pass filtered (25–250 Hz, 2-pass, 2nd-order Butterworth) and full-wave rectified. The TRI EMG data were normalized to its grand-mean of EMG activity for the last 200ms prior to the presentation of the goal target. We defined the spinal stretch reflex as the integrated EMG activity from 25-50ms following perturbation onset.

For *Experiment 1*, we used a one-way repeated measures ANOVA to compare the mean change in elbow angle and the elbow’s rotational velocity 20ms after perturbation onset, as well as the spinal stretch reflex from the TRI, as a function of how the robot applied mechanical perturbations at movement onset (3 levels: elbow flexed & wrist flexed; elbow flexed & wrist not perturbed; elbow flexed & wrist extended). Significant main effects were decomposed with planned paired sample t-tests.

For *Experiment 2*, we used a paired-sample t-test to compare the mean spinal stretch reflex from the TRI between the two blocks of trials (i.e., Upright vs Flipped) when the elbow and wrist were flexed, and when the elbow was flexed and the wrist was extended. All experimental results were considered reliably different if p <0.05.

## Results

### Experiment 1

There exists a spinal feedback pathway that integrates sensory information from the elbow and wrist and generates a spinal stretch reflex at elbow muscles that supports postural hand control (Weiler et al., 2019). Here we tested whether this putative spinal pathway can also efficiently assist in a reaching action. To do this, participants reached towards a target that required 10 degrees of elbow extension and we occasionally applied mechanical perturbations shortly after the participants began reaching. Perturbations always flexed the elbow, and simultaneously flexed the wrist, were not applied to the wrist, or extended the wrist.

### Features of behaviour

**Figure 1a,b** shows changes in elbow and wrist kinematics during trials in which mechanical perturbations flexed the elbow and wrist (green traces), only flexed the elbow (red traces), flexed the elbow and extended the wrist (blue traces) as well as during control trials, in which no perturbations were applied (grey traces). Note that we always applied the same 2Nm flexion perturbation to the elbow, but the elbow rotated differently as a function of the perturbation applied to the wrist. This was expected, as rotational forces applied or generated at one joint (e.g., the wrist) cause rotational forces at adjacent joints (e.g., the elbow). To highlight how quickly this relationship emerged we submitted the mean change in elbow angle 20 ms after perturbation onset when the wrist was flexed, not perturbed, and extended, to a repeated measures one-way ANOVA. Results of this analysis showed a reliable effect, F(2,38) = 270.0, p<0.001. Planned comparisons showed that the elbow angle when the wrist was mechanically flexed compared to when the wrist was not perturbed was reliably different, t(19) =7.8, p<0.001, and that the change in elbow angle when the wrist was not perturbed compared to when the wrist was mechanically extended was also reliably different, t(19) =21.3, p<0.001 (**Fig. 1d**).

**Figure 1.**
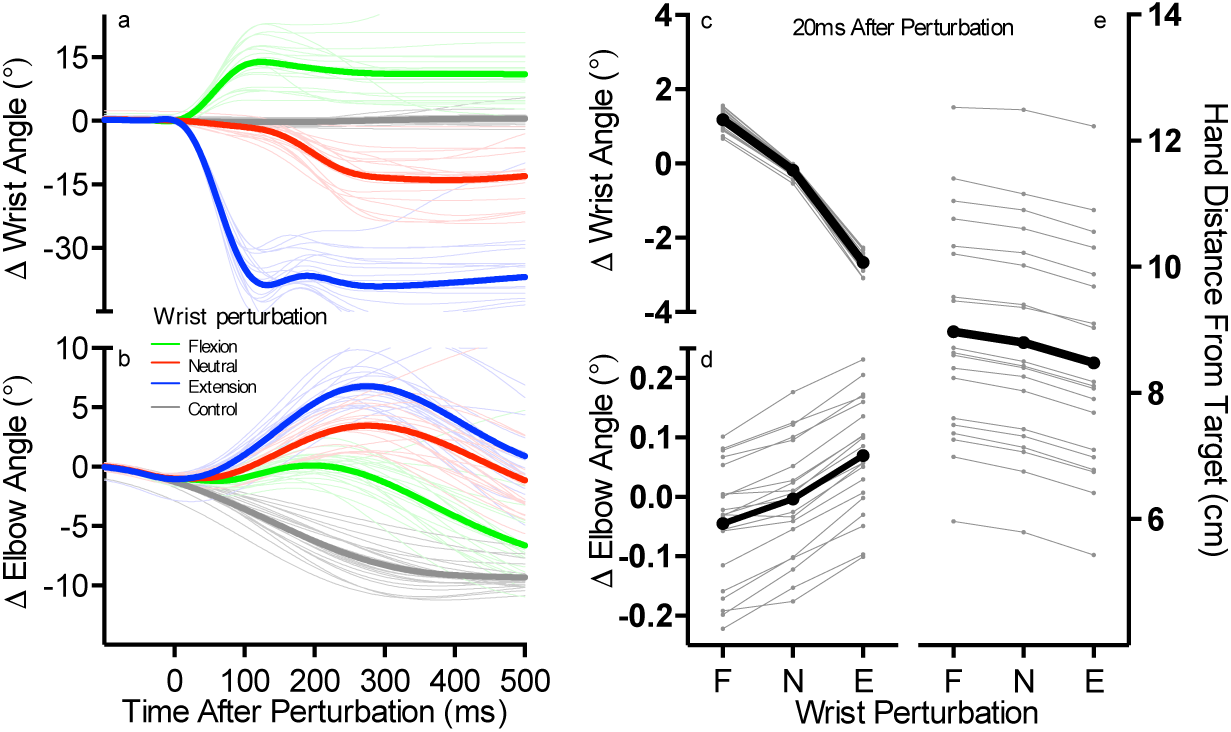
**a**: Change in wrist angle when the elbow was mechanically flexed and the wrist was simultaneously flexed (green), not mechanically perturbed (red), or extended (blue) at movement onset, and in control trials in which no perturbations were applied (grey). Data aligned to perturbation onset. Thin lines reflect individual participants and thick lines reflect the group mean. **b**: Same format as *a*, but for change in elbow angle. **c**: Mean change in wrist angle 20ms after perturbation onset for trials in which the elbow was flexed and the wrist was simultaneously flexed (F), not perturbed (N) and extended (E) at movement onset. Thin grey lines reflect individual participants and the thick black line reflect the group mean. **d**: Same format as *c* but for change in elbow angle. **e**: Same format as *c* but for the hand’s distance away from the target.

We then tested for differences in the elbow’s instantaneous rotational velocities 20ms after perturbation onset, as this provides insight to the state of the triceps (TRI) muscle (i.e., shortening, static, lengthening) in a temporal epoch that could causally influence the TRI spinal stretch reflex. To do this we computed the elbow’s rotational velocity at 20ms following the elbow flexion perturbation when the wrist was concurrently flexed, not perturbed, and extended, and compared each of these values to 0 deg/sec with single sample t-tests. These analyses showed that at 20ms post perturbation, the elbow was still extending when the mechanical perturbation flexed the elbow and wrist, t(19) = 2.70, p = 0.014, that the elbow was static when the mechanical perturbation only flexed the elbow, t(19) = 0.69, p = 0.49, and that the elbow had indeed begun to flex when the mechanical perturbation flexed the elbow and extended the wrist, t(19) = −7.71, p<0.001.

**Figure 1e** shows the hand’s mean distance from the target 20ms after perturbations that flexed the elbow and wrist, only flexed the elbow, and flexed the elbow and extended the wrist. We submitted these data to a repeated measures one-way ANOVA and results of this analysis also showed a reliable effect, F(2,38) = 918, p<0.001. Planned comparisons showed that the hand was further from the target when the wrist was flexed compared to when the wrist remain unperturbed, t(19) = 14.9, p<0.001, and that the hand was closer to the target when the wrist was extended compared to when the wrist remain unperturbed t(19) = −43.4, p<0.001.

### The elbow’s spinal stretch reflex

The kinematic analyses above shows that at 20ms after perturbation onset, the hand was furthest from the target when the elbow was still extending, whereas the hand was closest to the target when the elbow was flexing. The critical question here is whether the TRI spinal stretch reflex is tuned to how the elbow was rotating – and therefore how the TRI is shortening/lengthening – or is tuned to the hand’s distance from the target. If the former is true, the TRI spinal stretch reflex should be largest when the elbow was flexed the most by the perturbation (i.e., when the wrist was concurrently extended) and smallest when the elbow was flexed the least (i.e., when the wrist was concurrently flexed). In contrast, if the latter is true, the TRI spinal stretch reflex should be largest when the perturbation moved the hand furthest from the target (i.e., when the wrist was concurrently flexed) and smallest when the perturbation moved the hand close to the target (i.e., when the wrist was concurrently extended).

**Figure 2a** shows mean TRI EMG from trials in which mechanical perturbations flexed the elbow and wrist (green traces), only flexed the elbow (red traces), flexed the elbow and extended the wrist (blue traces) at movement onset, as well as during control trials, in which no perturbations were applied (grey traces). We submitted the mean EMG of the TRI spinal stretch reflex from trials that included mechanical perturbations to a repeated measures one-way ANOVA and results of this analysis showed a reliable effect, F(2,38) = 92.49, p<0.001 (**Fig. 2b**). Planned comparison showed that the TRI spinal stretch reflex was larger when the wrist was flexed compared to when the wrist was not perturbed, t(19) = 6.84, p<0.001, and that the TRI spinal stretch reflex was smaller when the wrist was flexed compared to when the wrist was not perturbed, t(19) = −10.13, p<0.001. This finding is consistent with a spinal feedback pathway that integrates information from the wrist and elbow to efficiently control hand movement.

**Figure 2.**
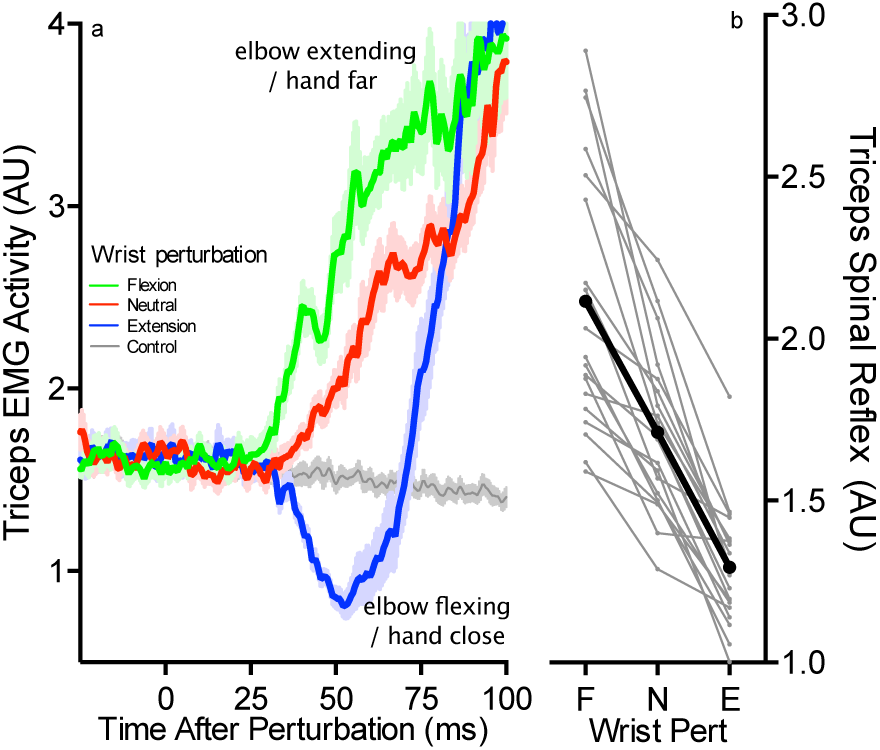
**a**: Mean EMG of the TRI when the elbow was mechanically flexed and the wrist was simultaneously flexed (green), not mechanically perturbed (red), extended (blue) at movement onset, and in control trials in which no perturbations were applied (grey). Data aligned to perturbation onset. Shading reflects ±1 SEM. Small inset panel shows mean EMG of the TRI across an expanded time range. **b**: Mean TRI spinal stretch reflex when the elbow was flexed and the wrist was simultaneously flexed (F), not perturbed (N) and extended (E) at movement onset. Thin grey lines reflect individual participants and the thick black line reflect the group mean.

### Experiment 2

In this experiment we tested whether the spinal feedback pathway can flexibly change its motor output to efficiently move the hand to a goal location when we diametrically altered how the rotation of the wrist moves the hand in external coordinates. This manipulation is accounted for during postural hand control (Weiler et al., 2019), but may not be present during reaching as there are notable architectural differences in the neural pathways that support remaining still compared to moving (Robinson, 1973, 1970; Shadmehr, 2017). To address this question participants again reached towards a target that required 10 degrees of elbow extension and we occasionally applied elbow flexion perturbations and simultaneous wrist flexion or extension perturbations at movement onset. Critically, participants completed these reaches in one block of trials by grasping the robot handle with their thumb pointing upward (i.e., Upright) and in another block of trials by internally rotating their forearm and grasping the handle with the thumb pointing downward (i.e., Flipped: **Figure 3**)

**Figure 3.**
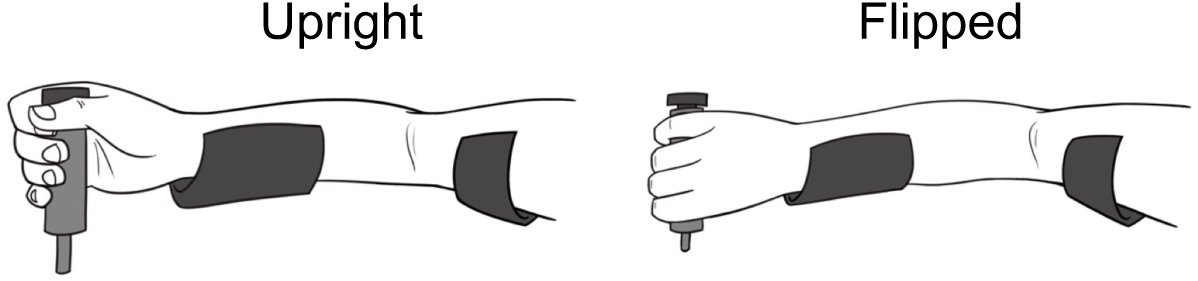
Cartoon of the Upright and Flipped arm orientations.

### Features of behaviour

Figure 4 shows how the Upright and Flipped arm orientations influenced the rotations of the elbow and wrist and movement of the hand when mechanical perturbations were applied at movement onset. There are two critical features of the data that we wish to highlight. First, changing the arm’s orientation influenced how the same mechanical perturbations moved the hand relative to the reaching target, and this was evident shortly after perturbation onset. Specifically, the hand was moved closer to the target 20ms after the elbow was flexed and the wrist was flexed when participants adopted the Flipped compared to the Upright orientation, t(11) = 9.32, p<0.001 (**Fig. 4e**). Conversely, the hand was moved closer to the target 20ms after the elbow was flexed but the wrist was extended when participants adopted the Upright compared to the Flipped orientation, t(11) = 10.65, p<0.001 (**Fig. 4j**). The second feature was that changing the arm’s orientation influenced how the elbow was flexed by the same mechanical perturbations, which was also evident 20ms after the perturbations were applied. Specifically, when the elbow and wrist were mechanically flexed, the elbow was flexed to a larger extent when participants adopted the Flipped compared to Upright orientation, t(11) = 9.07, p <0.001 (**Fig. 4b,d**). In contrast, when the elbow was mechanically flexed but the wrist was mechanically extended, the elbow was flexed to a larger extent when participants adopted the Upright compared to the Flipped orientation, t(11) = 2.74, p =0.019 (**Fig. 4g,i**). These two features in conjunction show that across both arm orientations, perturbations that moved the hand away from the target stretched the triceps the least, whereas perturbations that moved the hand close to the target stretched the triceps the most.

**Figure 4.**
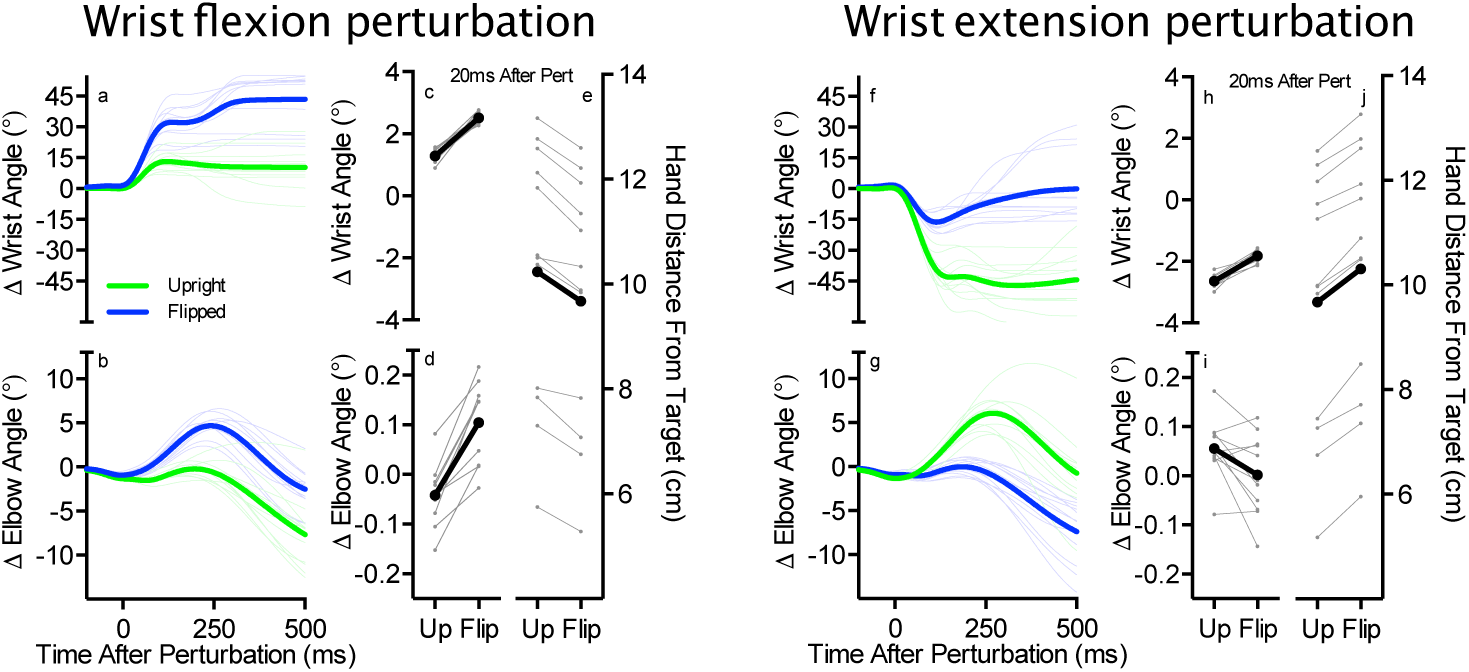
**a**: Change in wrist angle when the elbow and wrist were mechanically flexed at movement onset as a function of the Upright (green traces) and Flipped arm orientations. Thin lines reflect individual participants and the thick lines reflect the group mean. Data aligned to perturbation onset. **b**: Same format as A but for change in elbow angle. **c**: Change in wrist angle 20 ms after the elbow and wrist were mechanically flexed at movement onset for the Upright and Flipped orientations. Thin grey lines reflect individual participants and the thick black line reflect the group mean. **d**: Same format as C but for change in elbow angle. **e**: Same format as C but for the hand’s distance from the target. **F-J**: Same format as A-E but for when the wrist was mechanically extended.

We also note that the same wrist perturbations resulted in different amounts of wrist rotation as a function of the participants adopted arm orientation. This is not surprising as changing the arm orientation diametrically altered how the wrist should be rotated to move the hand to the target (**Fig a,f**). What was unexpected, however, was how quickly these differences manifested. Specifically, mechanical perturbations that flexed the wrist resulted in more wrist flexion 20ms following perturbation onset when participants adopted the Flipped compared to the Upright orientation, t(11) = 60.5, p<0.001 (**Fig. 3c**). In contrast, perturbations that extended the wrist caused more wrist extension 20ms following perturbation onset when participants adopted the Upright compared to the Flipped orientation, t(11) = 61.7, p<0.001 (**Fig. 4h**). These differences occur too quickly to be attributed to differences in volitional use of the wrist, and are thus most likely explained by slightly altered wrist biomechanics associated with grasping the robot manipulandum in different arm orientations. Regardless of the explanation, these differences also result in different rates in how the wrist muscles were stretched by the same wrist perturbations, and is a feature of the data we will return to in the Discussion.

### The elbow’s spinal stretch reflex

The kinematic analyses from this experiment showed that at 20 ms after perturbation onset, wrist flexion perturbations moved the hand closer to the reaching target but with a large amount of elbow flexion when participants adopted Flipped orientation, but moved the hand further from the reaching target with relatively little (if any) elbow flexion when participants adopted the Upright orientation. The reciprocal pattern was observed for wrist extension perturbations. The critical question here is whether the TRI spinal stretch reflex accounts for how changing the arm’s orientation influences the hand’s movement relative to the goal target.

**Figure 5a,b** shows mean TRI EMG from trials in which the elbow and wrist were flexed (top panel), and when the elbow was flexed and the wrist was extended at movement onset (bottom panel) as a function of the Upright (green traces) and Flipped (blue traces) arm orientation. We compared the TRI spinal stretch reflex across the two arm orientations and found that when the wrist was flexed, the TRI spinal stretch was larger when participants adopted the Upright compared to the Flipped arm orientation, t(11) = 5.98, p<0.001 (**Fig. 5c**). We also found that the when the wrist was extended, the TRI spinal stretch was larger when participants adopted the Flipped compared to the Upright orientation, t(11) = 9.57, p<0.001 (**Fig. 5d**). These findings are consistent with a spinal feedback pathway that can flexibly change its motor output to efficiently control hand movement.

**Figure 5.**
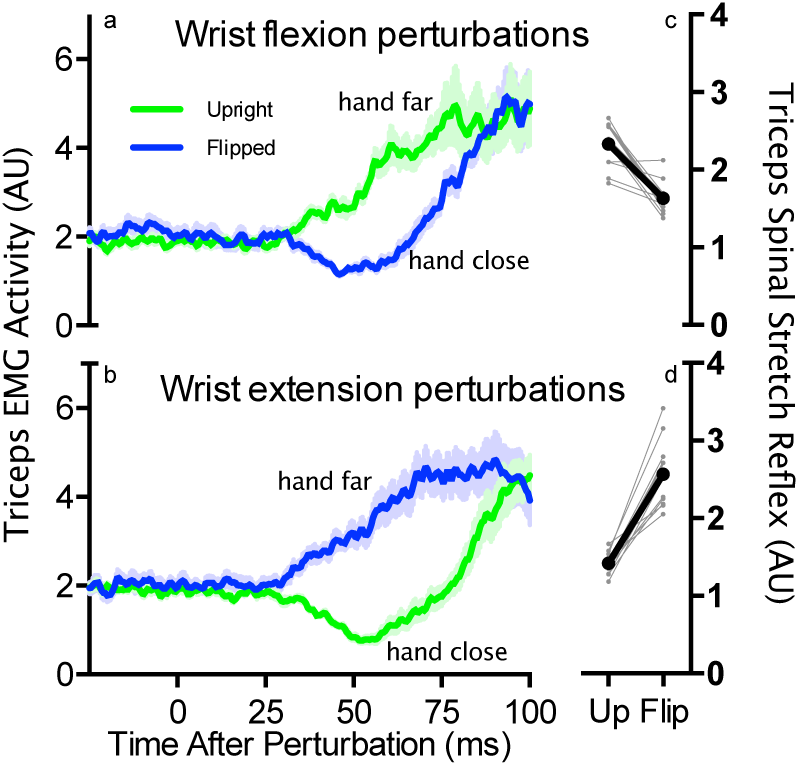
**a**: Mean EMG of the TRI when the elbow and wrist were mechanically flexed at movement onset when participants adopted the Upright (green traces) and Flipped (blue tracts) arm orientations. Data aligned to perturbation onset. Shading reflects ±1 SEM. **b**: Same format as A but for when the wrist was mechanically extended. **c**: Mean TRI spinal stretch reflex when the elbow and wrist were mechanically flexed at movement onset across the Upright and Flipped arm orientations. Thin grey lines reflect individual participants and the thick black line reflect the group mean. **d**: Same format as *c* but for when the wrist was mechanically extended.

## Discussion

In this study we applied mechanical perturbations that flexed the participant’s elbow and either flexed or extended their wrist once they began reaching towards a visual target. Critically, the perturbations that moved the hand towards the target did so with the most amount of elbow flexion, causing the most amount of TRI stretch, and the perturbations that moved the hand away from the target did so with the least (if any) amount of elbow flexion, causing little (if any) TRI stretch. Our results clearly show that TRI spinal stretch was not merely dependent on the amount the elbow was flexed – and thus the amount the TRI was stretched – but rather was tuned to the hand’s distance from the reaching target. We then had participants change the orientation of the arm, diametrically altering how the same rotation of the wrist moved the hand relative to the reaching target. When we mechanically perturbed the wrist and elbow in this new orientation, the TRI spinal stretch was also diametrically changed, and in such a way that was again tuned to the hand’s distance from the target. These findings are consistent with our previous work in which we showed that spinal stretch reflexes evoked in elbow muscles are tuned to the hand’s distance from its pre-perturbation static position (Weiler et al., 2019), indicating that the reflexes evoked to efficiently control the hand while reaching and while keeping the hand stationary are mediated by the same spinal circuity.

### Spinal processing for efficient hand control while reaching

Our movements are most efficiently controlled not by regulating the angular position of individual joints, but by generating corrective responses to the degree the overall goal of the action is hindered (Todorov, 2004; Todorov and Jordan, 2002). Implementing this control scheme requires understanding how multiple joints can be coordinated to move an end-effector (e.g., the hand), a level of sophistication thought to be an exclusive feature of the processing within a transcortical pathway that traverses areas critical for volitional movement production (Pruszynski and Scott, 2012; Scott, 2016). The notion that spinal circuits are incapable of such sophisticated is supported by the fact that the spinal circuity mediating the spinal stretch reflex, usually described as homonymous connections between muscle spindle and the spinal motor neurons of that same muscle (Pierrot-Deseilligny and Burke, 2012), often acts to regulate the position of individual joints independent of task constraints. Our first experiment showed that joint-regulation is not the functional limit of this putative spinal circuit. Specifically, this experiment clearly demonstrated that the circuit mediating the TRI stretch reflex was not acting to regulate or re-establish the angular position of the elbow, but rather was generating motor output that was tuned to the hand’s distance from the goal target. Generating this efficient motor output requires integrating sensory feedback from both the elbow and wrist, and there are documented heteronomous connections between the afferents of wrist muscles and motor neurons that innervate elbow muscles (Cavallari and Katz, 1989; Manning and Bawa, 2011; Weiler et al., 2019).

Our second experiment showed that these heteronomous connections from the wrist are not hardwired to the motor neurons that innervate the TRI. If that was the case, the TRI spinal stretch reflex would have been counterproductive when participants adopted the Flipped orientation. That is, when in the Flipped orientation, the TRI stretch reflex would have been largest when the hand was closest to the target and inhibited relative to baseline levels of muscle active when the hand was far from the target. Recall also that wrist perturbations flexed the wrist more in the Flipped compared the Upright orientation and extended the wrist more in the Upright compared the Flipped orientation (**Fig. 4c,h**). If the heteronomous wrist-to-TRI connections were indeed hardwired, we would have expected larger TRI spinal reflexes for the Flipped compared the Upright orientation for trials when the wrist was mechanically flexed and larger spinal reflexes in the TRI for the Upright compared to the Flipped orientation trials when the wrist was mechanically extended. But this is not what we observed. Instead, we observed a reversal of the TRI spinal stretch reflex when participants changed the orientation of the arm, which was again tuned to the hand’s distance from the reaching target. This indicates that this spinal pathway has a mechanism that takes into account the arm’s orientation to selectively gate what afferent information from the wrist is allowed to influence the TRI motoneuron pool. This may be accomplished through presynaptic inhibition which, based on the arm’s orientation, regulates whether the stretched wrist muscle provides the triceps’ motorneuron excitatory input through a monosynaptic direct pathway or inhibitory input via an indirect pathway routed through an inhibitory interneuron (**Fig. 6**; see also Seki et al., 2003). We have already determined it is the wrist extensor muscles that influence the motorneurons that innervate the triceps whereas the wrist flexor muscles influence the motorneurons that innervate the biceps (Weiler et al., 2019). Future work using animal models will be needed to determine if the presynaptic mechanism proposed here exists and whether it is governed by spinal processing that locally determines the arm’s orientation or is implemented by supraspinal regions of the nervous system.

**Figure 6.**
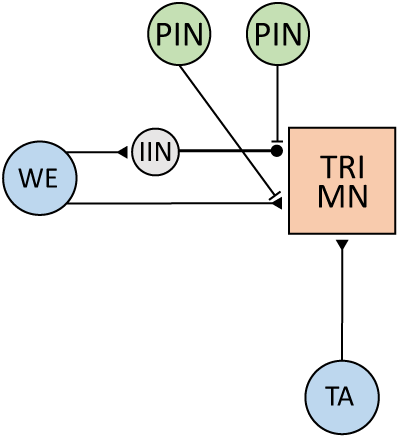
Cartoon of proposed spinal circuit. The stretched triceps muscle (TA) and the stretched wrist muscle (WA) provide excitatory input into the triceps’ spinal motor neurons (TRI MN). Inhibitory input into the TRI MN is also provided from the stretched wrist extensor muscles (WE) through an inhibitory interneuron (IIN). Separate pools of presynaptic inhibitory neurons (PIN) selectively gate whether the excitatory or inhibitory input from the wrist muscle exert their influence on the triceps’ motor neurons as a function of the arm’s orientation.

### Functional versus volitional control

A key distinction between spinal and supraspinal feedback pathways is the degree to which they can convert sensory information to motor output; the output from spinal pathways are rapidly generated in a relatively simple fashion, being tightly coupled to the sensory input and the current state of the spinal circuity, whereas the output from a supraspinal pathways are generated in a more deliberate and sophisticated manner, reflecting seemingly any volitional sensory-to-motor transformation (for review, see (Pruszynski and Scott, 2012). We highlight this distinction because we have shown that a spinal pathway efficiently supports the volitional act of reaching, but we do not intend to imply that the output of this pathway is dependent on an individual’s volitional intention to reach. In fact, when individuals are instructed to “not intervene” following simultaneous elbow and wrist perturbations, a spinal pathway generates an efficient response that attempts to return to the hand to its original position even though this is against the individual’s intent (Weiler et al., 2019). This is not to say that individuals are incapable of volitionally tuning spinal reflexes. Indeed, with weeks of training, the H-reflex can be up- or downregulated (Thompson et al., 2013; Wolpaw, 1980), which may provide a novel therapeutic approach for treating spasticity following spinal cord injury. However, immediately modulating the output of spinal pathways by volitional intent appears to be a common limitation of how these neural circuit process sensory information.

Despite the inability to volitionally modulate the immediate motor output from spinal pathways in real-time, the spinal reflexes we showed here and previously – which help to efficiently move to a new location or keep the hand stationary – are functional for how we use our hands in the real-world. This is similar to the vestibulo-ocular reflex, which is another rapid response that is refractory to an individual’s volitional intent yet functionally relevant for stabilizing the visual inputs on the retina during real-world events. Perhaps instead of contrasting spinal and supraspinal pathways by the apparent sophistication in which they can process sensory information, a better contrast may relate to how flexible this processing can be. On the one hand supraspinal pathways can transform sensory inputs into motor responses in a seemingly arbitrary way, whereas on the other hand spinal pathways process sensory feedback in a very regulated fashion and in a way that supports specific core functions of particular ethological importance.

### Sensory gating and movement control

A primary motivator for this work was that sensory information can be attenuated at various levels of the nervous system when the body is in motion (Ghez and Pisa, 1972; Hantman and Jessell, 2010; Papakostopoulos et al., 1975; Seki and Fetz, 2012; Tsumoto et al., 1975). This effect is typically associated with changes in conscious percepts. For example, active movement increases the threshold to perceive a sensory stimulus (Chapman et al., 1987; Duysens et al., 1990) and hinders a person’s ability to discriminate between two sensory stimuli (Conte et al., 2016). This sensory gating mechanism and its influence on perception may seem counterproductive from a movement control point of view because the act of generating accurate movements requires the constant processing of sensory information (Rothwell et al., 1982). How does the movement-induced suppression of sensory feedback not disrupt the execution of purposeful movement? We propose two potential mechanisms.

First, is that sensory-gating may be restricted to neural circuity involved in the generation of percepts, leaving sensory feedback used for real-time movement control unfettered. Such an idea may be best supported by seminal work that investigated how saccadic suppression influences reaching, which showed that individuals are able to quickly alter their reaching trajectory when a visual target changes position during a saccade even though the target-jump was imperceptible (Goodale et al., 1986). And second, is that sensory inputs may not be gated if such information governs aspects of the movement. Indeed, the ability to detect a vibrotactile stimulus on the forearm is reduced during a reaching and grasping action compared to when at rest. When the same stimulus is applied to the fingers of the grasping hand, detection thresholds are unaffected (Colino et al., 2014). Similarly, tactile processing of the geometric features of touched objects is nearly an order of magnitude more precise when a person is engaged in an object manipulation task than when they are performing a perceptual judgement (Pruszynski et al., 2018).

